# Genome-centric metagenomics revealed the spatial distribution and the diverse metabolic functions of lignocellulose degrading uncultured bacteria

**DOI:** 10.1101/328989

**Authors:** Panagiotis G. Kougias, Stefano Campanaro, Laura Treu, Panagiotis Tsapekos, Andrea Armani, Irini Angelidaki

## Abstract

The mechanisms by which specific anaerobic microorganisms remain firmly attached to lignocellulosic material allowing them to efficiently decompose the organic matter are far to be elucidated. To circumvent this issue, the microbiomes collected from anaerobic digesters treating pig manure and meadow grass were fractionated to separate the planktonic microbes from those adhered to lignocellulosic substrate. Assembly of shotgun reads followed by binning process recovered 151 population genomes, 80 out of which were completely new and were not previously deposited in any database. Genome coverage allowed the identification of microbial spatial distribution into the engineered ecosystem. Moreover, a composite bioinformatic analysis using multiple databases for functional annotation revealed that uncultured members of *Bacteroidetes* and *Firmicutes* follow diverse metabolic strategies for polysaccharide degradation. The structure of cellulosome in *Firmicutes* can vary depending on the number and functional roles of carbohydrate-binding modules. On contrary, members of *Bacteroidetes* are able to adhere and degrade lignocellulose due to the presence of multiple carbohydrate-binding family 6 modules in beta-xylosidase and endoglucanase proteins or S-layer homology modules in unknown proteins. This study combines the concept of variability in spatial distribution with genome-centric metagenomics allowing a functional and taxonomical exploration of the biogas microbiome.

**Importance:** This work contributes new knowledge about lignocellulose degradation in engineered ecosystems. Specifically, the combination of the spatial distribution of uncultured microbes with genome-centric metagenomics provides novel insights into the metabolic properties of planktonic and firmly attached to plant biomass bacteria. Moreover, the knowledge obtained in this study enabled us to understand the diverse metabolic strategies for polysaccharide degradation in different species of *Bacteroidetes* and *Clostridiales*. Even though structural elements of cellulosome were restricted to *Clostridiales*, our study identified in *Bacteroidetes* a putative mechanism for biomass decomposition based on a gene cluster responsible for cellulose degradation, disaccharide cleavage to glucose and transport to cytoplasm.

## Introduction

The mechanism of several microbial processes remains unclear due to the fact that only a small fraction of microorganisms that exists in nature can be cultivated (1). Especially in complex microcosms, such as the one responsible for anaerobic digestion of organic matter, the understanding of the microbial dynamicity is additionally hampered by the presence of syntrophic interactions between members of the community. During the past years, different molecular techniques, ranging from 16S rRNA gene sequencing to metagenomics, were proposed to overcome these obstacles and to explore the functional principles of the unculturable microbial majority.

In comparison to 16S rRNA gene based studies, metagenomic investigations using shotgun sequencing became more attractive and were applied to anaerobic digestion systems to obtain relevant insights on the metabolic properties of the whole microbiome (2). However, the reads generated by high-throughput sequencing are short, error-prone and contain limited signal for homology searches; thus, shotgun assembly rendered more reliable (3, 4). Moreover, all these “gene-centric” metagenomic studies consist of inventories of individual annotated genes (5), without classifying the genes to single population genomes (PG). Recent advances in bioinformatics and introduction of automated strategies for binning process, such as CONCOCT, GroopM, or MetaBAT (6–8), shift the perspective of metagenomics from “gene-centric” to “genome-centric”. Quite recently, this novel approach was applied to the anaerobic digestion microbiome leading to the extraction of hundreds of PGs, which were assigned to the four steps of the process, i.e. hydrolysis, acidogenesis, acetogenesis, and methanogenesis (9–11). The genome-centric metagenomics have the undoubted benefit of revealing the functional properties of individual genomes, leading to a more detailed comprehension of the microbial interactions occurring into the microbiome (12). Additionally, the identification of the sequenced genomes can be combined with metatranscriptomic data providing further information regarding the metabolic basis for adaptation of particular phylotypes (5) and allowing the monitoring of gene expression changes to single-species level (13).

The microbiomes associated with the conversion of lignocellulosic-derived sugars into biofuels and bioproducts are gaining particular attention in the wake of climate change. It is well known that the hydrolysis step of anaerobic lignocellulose digestion process is mainly facilitated by some anaerobic bacteria that are able to produce a multienzyme complex, called cellulosome, and by aerobic bacteria and fungi that typically produce free, non-complexed individual enzymes (14, 15). Moreover, the recalcitrant nature of the lignocellulosic substrate renders the hydrolysis process as a rate-limiting step in the anaerobic digestion food chain (16). Therefore, a more detailed knowledge on the firmly attached species and their interactions with the planktonic microbiome is essential for the development of strategies that will accelerate the hydrolysis rate. In turn, the overall efficiency of biotechnological processes (e.g. production of biofuels and bio-based platform chemicals) will be significantly improved.

Previous studies targeting the identification of the microbial species firmly attached to the plant material were based only on 16S rRNA gene-based terminal restriction fragment length polymorphism (17) or FISH (18). Recently, metagenomic binning analysis and metatranscriptomics were performed to identify species involved in cellulose degradation (10, 19). Nevertheless, these high throughput sequencing methods did not study the spatial distribution of the microbial species. Moreover, detailed genomic analyses on polysaccharides binding and degradation pathways were focused on few microbial species, with *Clostridium thermocellum* to be recognized as the most interesting model (20). However, limited information is available for uncultured species of *Bacteroidetes* phylum (21), which are known to cover a relevant role in proficient plant biomass-degrading ecosystems and in anaerobic digestion sludge (19, 22). The importance of *Bacteroidetes* in thermophilic cellulose methanation process was previously underlined (19), and therefore, the sequencing of these genomes is extremely important to shed light on their metabolic potential.

In the present study, a genome-centric approach was applied to elucidate the spatial distribution of lignocellulose degradation mechanism in anaerobic digestion process. This was done as it was hypothesised that the localisation of microorganisms should be mainly determined by their functional roles, and thus, the microbes that are firmly attached to the plant material should be responsible for the degradation of lignocellulosic biomass. Samples obtained from biogas reactors (i.e. firmly-attached to grass particles and planktonic phases of co-digestion system and independent manure samples from mono-digestion system) were subjected to Illumina high-throughput shotgun sequencing. Subsequent *de novo* assembly and binning process led to the extraction of 151 population genomes. Bioinformatics functional analysis performed on the recovered draft genomes allowed a detailed characterization of the functional structure of the microbiome, and more importantly, clarified the strategies followed by different species to bind the solid fraction of plant material and to perform polysaccharides hydrolysis.

## Results

A targeted fractionation strategy was performed to analyse the spatial localisation of the microbiome involved in lignocellulose degradation in biogas reactors. More specifically, nine samples were collected (i.e. three biological replicates per each treatment) representing the microbial consortia residing in the Liquid Grass fraction (LG), or Firmly attached to Grass biomass (FG), or in an anaerobic system treating Pig Manure without lignocellulose (PM). Approximately 120 million paired-end reads were co-assembled to obtain a reliable representation of the genomes present in the microbial communities. This process resulted in the generation of 483,935 scaffolds ranging in size from 500 to 371,582 bp (N50 2426 bp; total length 775,466,133 bp). Gene prediction performed on the scaffolds led to the identification of 1,102,319 protein-encoding sequences whose function were investigated using five software packages based on different databases (i.e. KEGG, COG, dbCAN, SEED, Pfam). The combination of diverse annotation strategies provided a detailed overview of the functional properties of the metagenome.

The applied binning process allowed the extraction of 151 PGs from the assembly (Fig. 1) with an estimated completeness ranging between ∽11% and 100% (average 73%) (Fig. 1 and Supplemental Dataset S1). The superlative outcome of the assembly and binning processes was evidenced by the 120 PGs and the 94 PGs having an estimated completeness of more than 50% and 70%, respectively. Moreover, 45 PGs fulfil the stringent criteria for defining “high quality draft” genomes (i.e >90% complete with less than 5% contamination) according to the recently published standard metrics (23). PGs with lower than 50% completeness were only used to define the taxonomy and to check species abundance in the three different fractions. Thus, the functional characteristics derived from their gene content are not included in the discussion section.

**Fig. 1.**
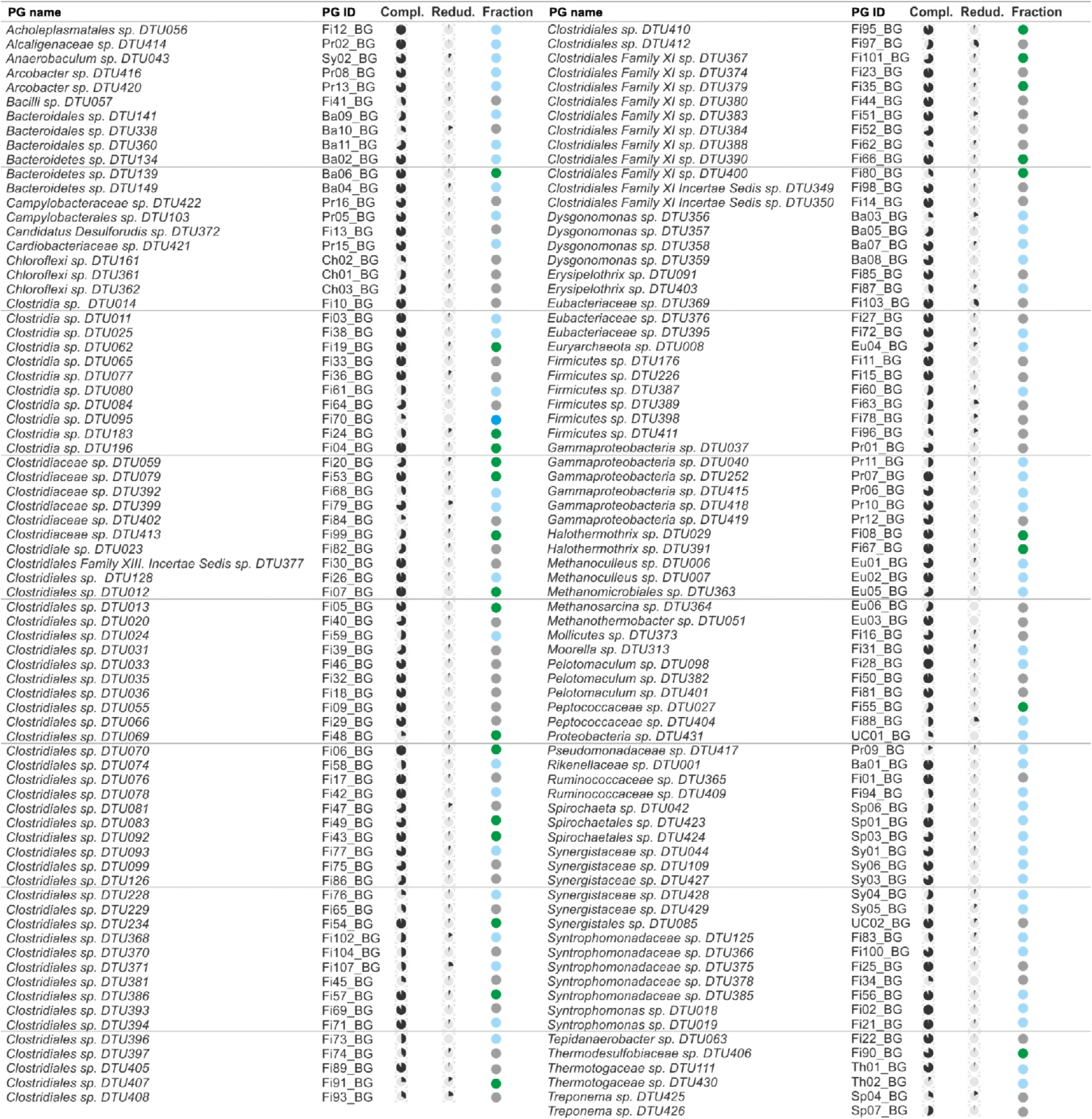
Characteristics of the recovered PGs. The columns designate: extended PG names, short IDs, pie graphs representing the completeness of the PGs, pie graphs representing their contamination level and a colour code indicating PG that were firmly attached to the grass (green), planktonic (light blue) and those that were similarly enriched in both fractions (gray).

The taxonomy structure of the biogas microbiome was dominated by members of *Firmicutes* (101 PGs), followed by *Bacteroidetes* (11 PGs), *Gammaproteobacteria* (8 PGs), *Synergistia* (7 PGs) and some other less frequently found phyla (Fig. 1 and Supplemental Dataset S1). Archaea were represented by 5 PGs; four of them belonging to *Methanomicrobia* and one to *Methanobacteria* (Fig. 1). All the extracted PGs were present in the three fractions (i.e. FG, LG and PM) in different coverage levels, except from one methanogen (Euryarchaeota sp. DTU008) that was absent from FG samples.

Analysis of the PGs coverage in the 9 examined samples revealed that most of the reconstructed genomes (64) were enriched more than 2 folds in samples collected from the liquid fraction (LG), while only 25 species were found to increase more than 2 folds in the samples collected from the grass biomass (FG) (Fig. 1 and Fig. 2). The remaining 62 PGs did not present any significant change in any of these two fractions (Fig. 2). Some fundamental functions, such as methanogenesis, are preferentially restricted to the liquid fraction as verified by *Methanomicrobiales* species (Eu01_BG, Eu02_BG, Eu05_BG) that were identified only to the planktonic microbiome (Fig. 1 and Fig. 3). Other archaeal species belonging to the orders *Methanobacteriales* (Eu03_BG) and *Methanosarcinales* (Eu06_BG) have an “intermediate behaviour” as they were identified both in FG and in the LG fractions. Relative abundance of bacteria and archaea in the two fractions was verified using real-time PCR with universal primers (27F, 1492R) and (109F, 1492R). Results obtained revealed a 2.1 fold enrichment of bacteria in the firmly attached to grass fraction and a 4.8 fold enrichment of archaea in the liquid fraction. A comparison between the gene contents of firmly attached to the grass PGs and the planktonic PGs were the initial stepping stone to identify the functional differences among these two groups. COG analysis showed that the number of genes in 9 clusters of orthologous groups of proteins was significantly different between the two microbial groups (Supplemental Dataset S1). Specifically, the firmly attached to grass PGs were more enriched in categories “G-Carbohydrate transport and metabolism”, “T-Signal transduction mechanisms” and “N-Cell motility”, while the planktonic PGs were more enriched in “C-Energy production and conversion”. The presence of numerous genes belonging to the planktonic PGs in COG category “C-Energy production and conversion” is related to their ability to use intermediate metabolites derived from plant polysaccharides degradation, such as volatile fatty acids, for energy production.

**Fig. 2.**
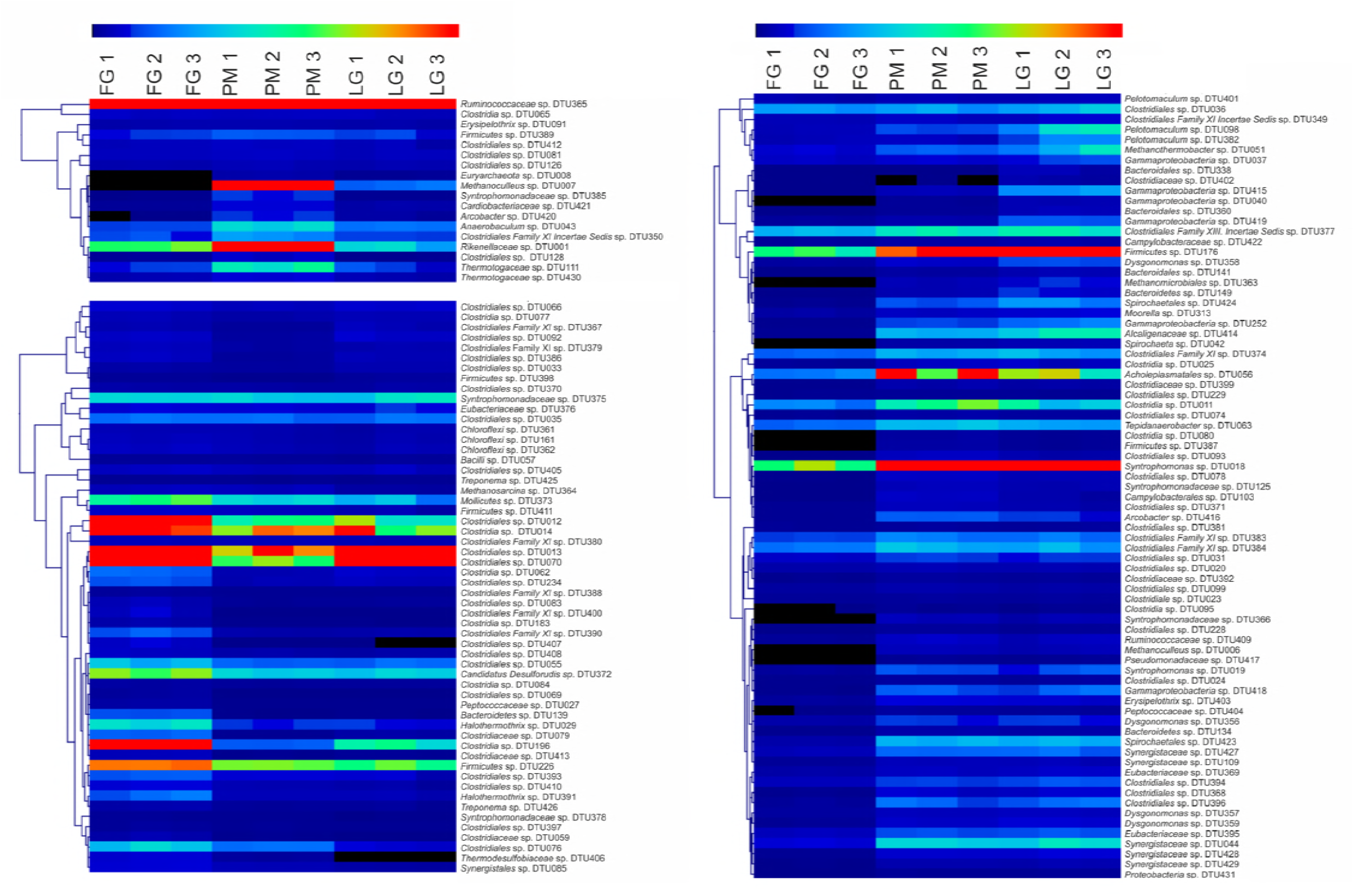
Abundance of the 151 PGs is represented as a heatmap. Three replicates were collected from the microbial communities which were firmly attached to the grass (FG), or residing the liquid phase of the reactor (LG), or populating the reactor treating exclusively pig manure (PG). For better visualisation of the outcomes, the heatmap was split into three sections based on the hierarchical clustering of the extracted PGs. Correspondence between colors and coverage profile is illustrated with a scale (i.e. decreasing coverage is ranking from red to black) depicted above the heatmaps.

**Fig. 3.**
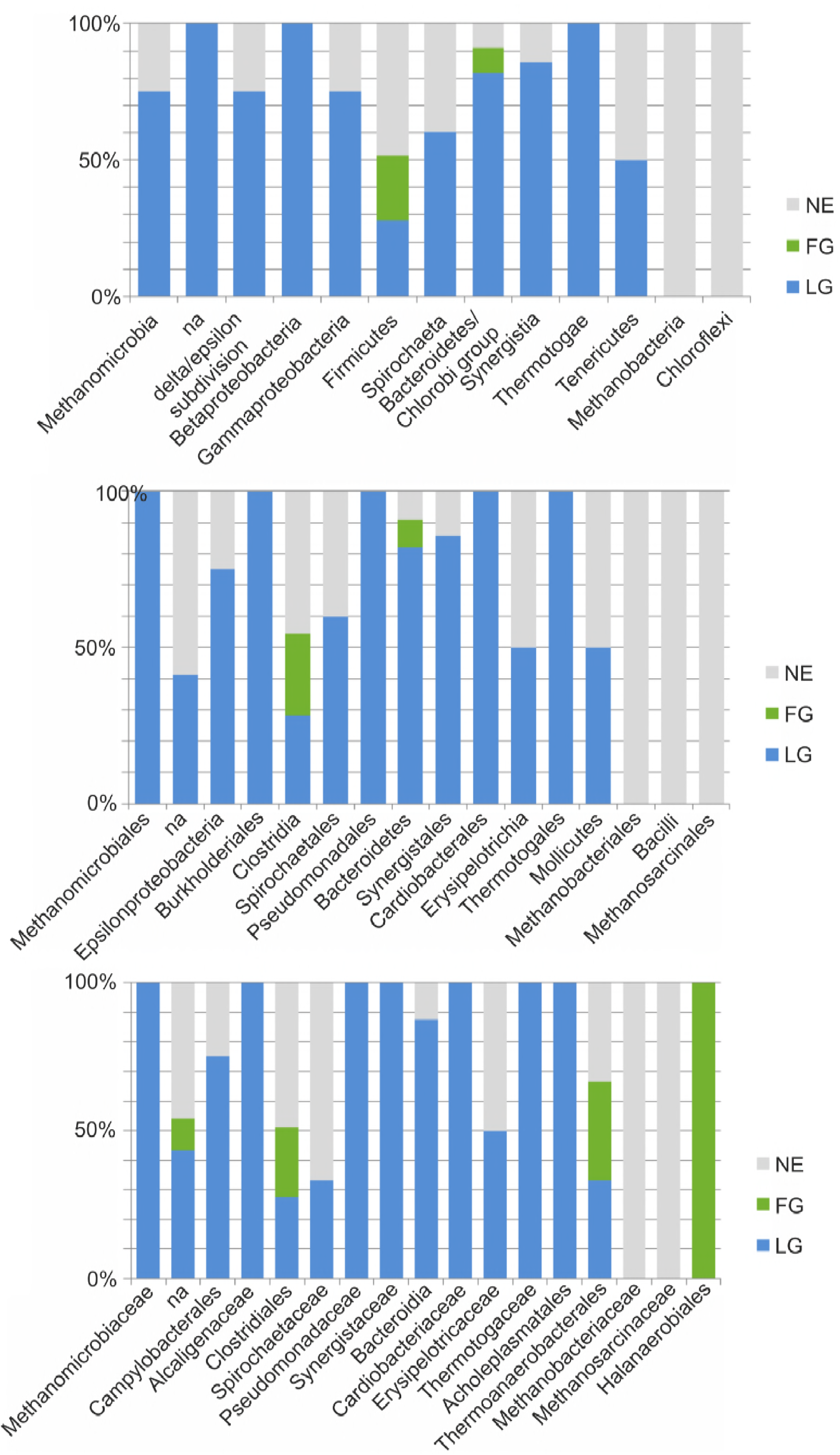
Comparative taxonomic analysis of the PGs. The PGs were grouped as firmly attached to the grass (green), planktonic (blue) and those that were similarly enriched in both fractions (gray). For each taxa, the proportion of PGs belonging to the three groups is represented. In (A) the PGs are classified at phylum level, in (B) at family level and in (C) at order level.

A deeper insight between the diverse metabolic properties of the firmly attached to grass and the planktonic PGs can be obtained from the SEED subsystem (second level annotation), which provides a more specific functional assignment of the proteins (Supplemental Dataset S1). This analysis confirmed that the firmly attached to grass PGs encode more proteins related to “motility” and “chemotaxis”, and additionally, revealed an enrichment in “adhesion” and “biofilm formation” categories. The latter was also supported by the presence in these PGs of a higher number of proteins associated to RNA polymerase sigma-54 factor rpoN, which is a pleiotropic transcription factor involved in regulation of genes related to carbohydrate metabolism, motility and biofilm formation (24). Moreover, the ability of the firmly attached to grass PGs to perform carbohydrate degradation and metabolism is proven by the presence of numerous genes involved in “di-and oligosaccharides” and / or “monosaccharides” utilization and “glycoside hydrolases”. The planktonic PGs encode on average a higher number of proteins involved in transport and utilization of simple carbon compounds derived from the degradation of complex plant polysaccharides such as “organic acids” and “tricarboxylate transporters”. Moreover, they contain higher number of proteins involved in “one-carbon metabolism”, “cofactors, vitamins, prosthetic groups, pigments” and “coenzyme F420” demonstrating a closer involvement during the process of methanogenesis.

The results from the dbCAN database (Supplemental Dataset S1) showed that the detected glycoside hydrolases families included known carbohydrate-active enzymes, which were classified in five categories, namely cellulases, endohemicellulases, debranching enzymes, oligosaccharide-degrading enzymes and other glycoside hydrolases. The PGs having the higher number of glycoside hydrolases domains for each category are illustrated in Fig. 4. The analysis demonstrated that the firmly attached to grass PGs have more cellulases (particularly *Clostridia* sp. DTU196, *Clostridiaceae* sp. DTU079 and *Clostridiales* sp. DTU092), while PGs in the liquid fraction have more debranching enzymes (particularly *Bacteroidetes* sp. DTU149, *Dysgonomonas* sp. DTU359 and *Clostridiales* sp. DTU078).

**Fig. 4.**
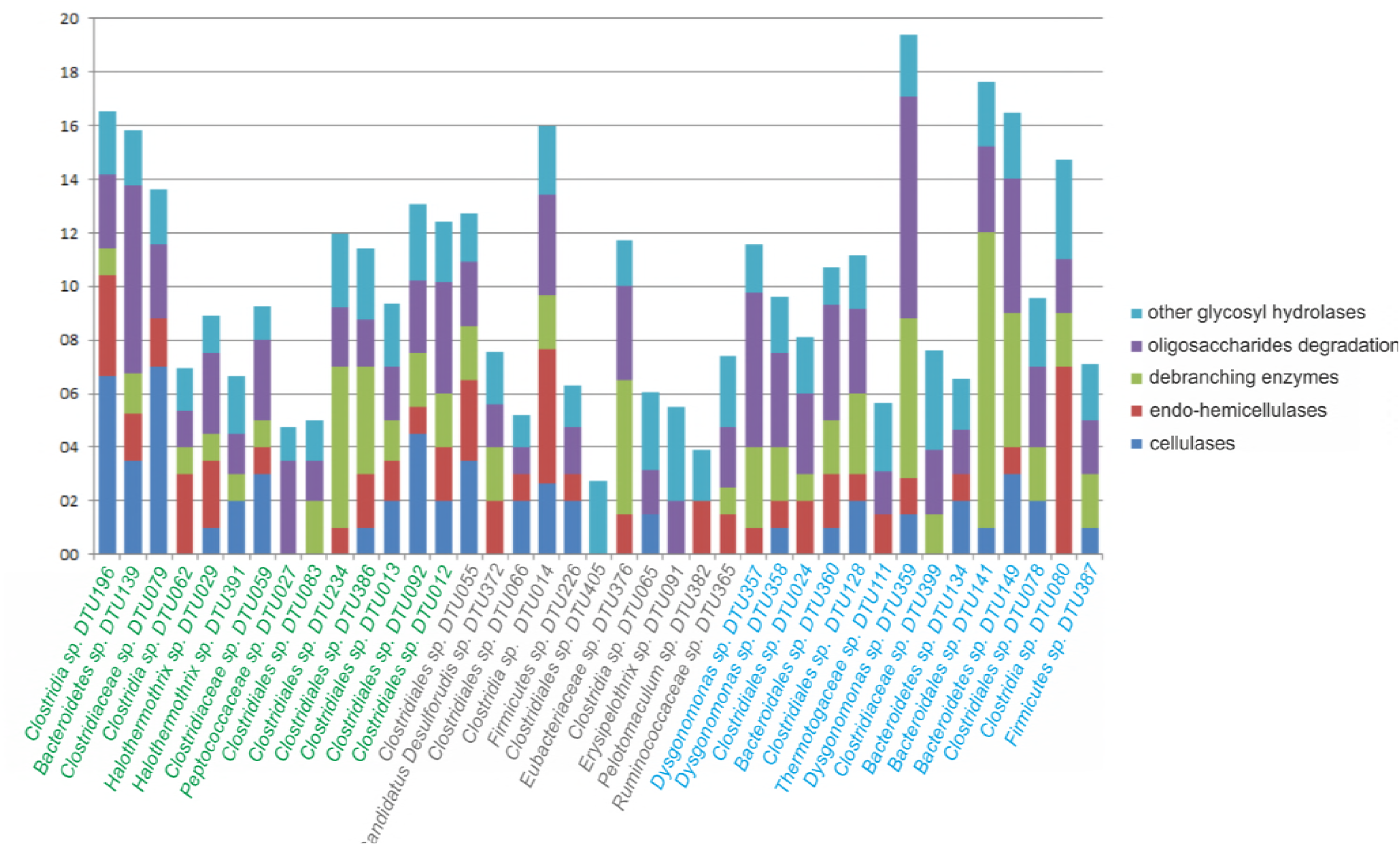
Number of carbohydrate active domains in PGs. PGs with the higher number of GH domains and with estimated completeness higher than 50% were analyzed. Domains belonging to the GH were grouped in four classes considering their role in carbohydrates degradation and the remaining GH are described as “other GH”. PGs were coloured depending on their spatial location, firmly attached “green”, “liquid fraction” blue, intermediate behaviour “gray”.

Among the nearly-complete population genomes, *Clostridiaceae sp.* DTU079, *Bacteroidetes sp.* DTU139 and *Clostridia sp.* DTU196 exhibited the highest enrichment in the FG samples (Supplemental Dataset S1). More specifically, the coverage profile of these PGs was found to be significantly increased in the FG samples compared to the LG by 14 (p<0.001), 19 (p<0.0003) and 25 (p<0.003) fold, respectively. Analysis of the carbohydrate binding modules among the reconstructed PGs evidenced that only the *Firmicutes* genomes *Clostridiaceae sp.* DTU079 and *Clostridia sp.* DTU196 contain numerous cohesin and dockerin domains. The cohesin-dockerin pairs constitute fundamental elements of cellulosome, which in turn can efficiently degrade plant polysaccharides (e.g. cellulose). The cellulosome structure of *Clostridiaceae sp.* DTU079 contains 12 cohesin and 61 dockerin domains (distributed on 6 and 36 proteins, respectively) and is much more complex in comparison to *Clostridia sp.* DTU196, which contains 12 cohesin and 26 dockerin domains (on 7 and 17 proteins, respectively) (Fig. 5; Supplemental Dataset S1). COG annotation of the *Clostridiaceae sp.* DTU079 cellulosome revealed that many of its enzymatic activities are represented by beta-xylosidases and endoglucanases, while the *Clostridia sp.* DTU196 cellulosome contains mainly beta-xylosidases. The evaluation of the genomic localization of genes encoding cellulosome components demonstrated that they are comprised in defined polysaccharides utilization loci. In *Clostridia sp.* DTU196 there are seven main regions localized on scaffolds 13, 48, 286, 790, 944, 1201 and 1476, while in *Clostridiaceae sp.* DTU079 there are nine main regions on scaffolds 531, 956, 1421, 4817, 4977, 7006, 8003, 8935 and 9186. Moreover, some other genes encoding subunits of the cellulosome are scattered along the genome.

**Fig. 5.**
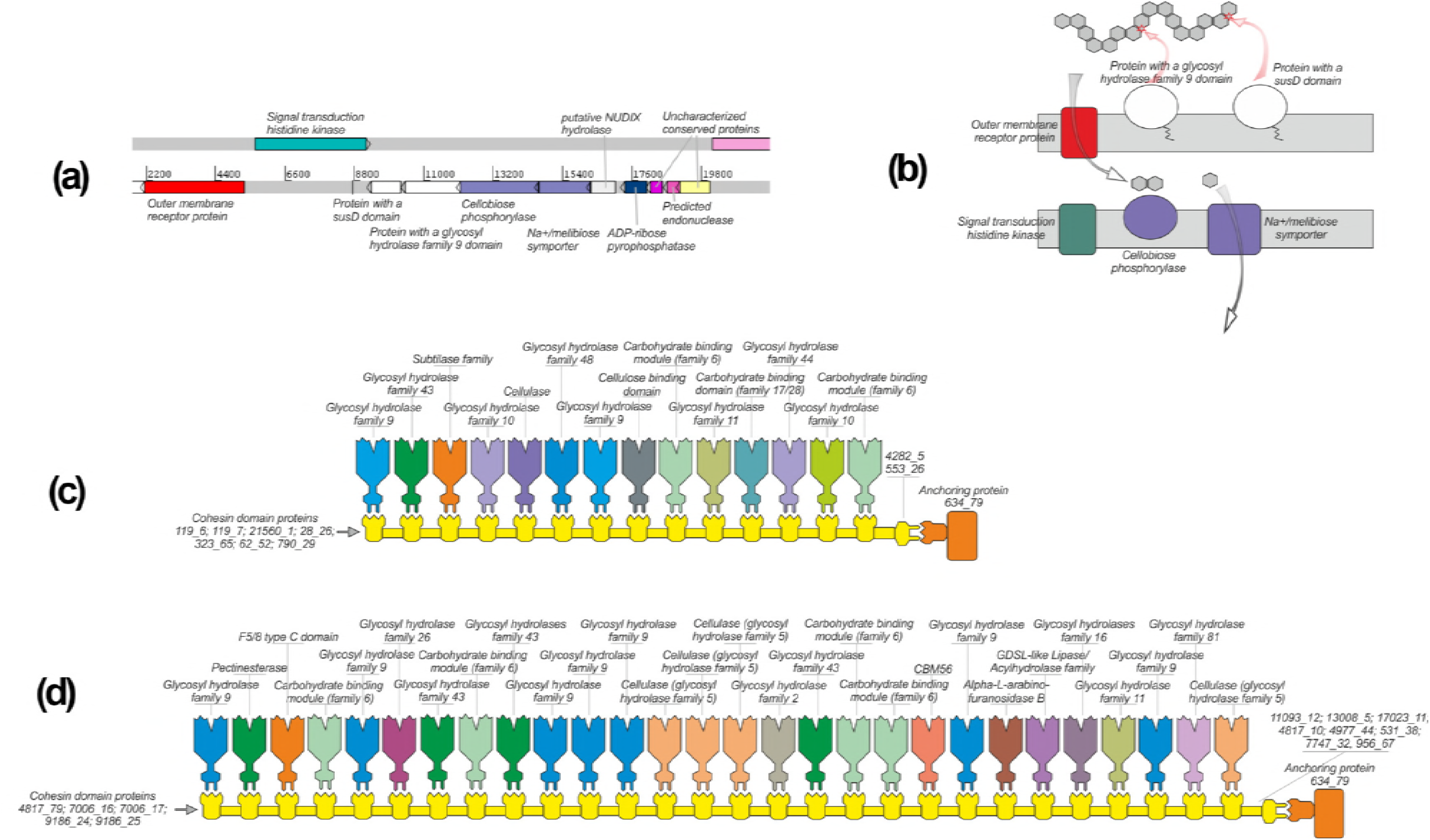
PULs organization and cellulosomes of Fi04_BG and Fi53_BG. (a) Gene organization of a relevant PUL in Ba06_BG. Genes are colored according to their COG function. (b) Proteins encoded by the genes putatively involved in cellulose degradation are assigned to their cellular localization and their putative role is reported according to the functional annotation and to the model proposed by Martens and coll.(2009). (c-d) Schematic representation of the cellulosome complex identified in Fi04 and Fi53. The putative functional role is reported according to their annotation.

On the contrary, the genes involved in the mechanism of polysaccharide utilization in the *Bacteroidetes* sp. DTU139 are clustered on six specific regions (i.e. on scaffolds 3625, 9343, 18620, 21571, 22665 and 22746). The most interesting polysaccharides utilization loci was found on scaffold 18620 and it is composed by 8 genes encoding respectively an outer membrane receptor protein (18620_3), a signal transduction histidine kinase (18620_4), two proteins of unknown function (18620_5, 18620_6), a cellobiose phosphorylase (18620_7) and a disaccharide transporter (18620_8) (Fig. 5a). The unknown protein “18620_5” contains a susD (Starch-binding protein) domain, while 18620_6 contains a glycosyl hydrolase family 9 domain.

## Discussion

The high completeness level of the recovered PGs allowed a detailed determination of the metabolic potential of the microbial population. The taxonomic assignment of the microbial members showed that the dominant phyla of the microbiome were in accordance with previous studies (2, 9, 25). It should be underlined that 80 out of the 151 recovered PGs were newly identified, while the rest PGs (i.e. 71 PGs, 47% of the total community) were the same species found in previous metagenomic binning studies (Fig. S1) (26, 27). This result clearly demonstrates that the engineered anaerobic digestion microcosm is far to be elucidated, and further investigations are mandatory to identify the numerous species that currently remain unknown at genome level. The newly identified PGs represented 53% of the planktonic microbiome and 40% of the community that was firmly attached to grass biomass. The relatively higher portion of newly identified PGs in the planktonic microbiome can be attributed to the utilisation of pig manure as co-substrate. As extensively described in the Supplemental Information S1, the diverse chemical composition of pig manure (mainly due to high ammonia concentration) compared to the cattle manure affected both the bacterial and archaeal community profile. The distinctively different microbial communities of biogas reactors treating cattle manure or pig manure are evidenced by plotting as heatmap the coverage profiles of their metagenomes (Supplemental Information S1). It has been previously reported that environmental variables, such as the physicochemical conditions of the feedstock, govern the microbial community structure and their linkage with the process efficiency in anaerobic digesters (28). The enrichment of specific microbes in the firmly attached to grass biomass could classify them as putatively responsible for lignocellulose degradation. More specifically, these microbes could bind to grass particles and fibres that are present in the reactor, and consequently, the proximity of the microorganisms with the substrate facilitates the plant cell wall deconstruction process. It was shown that only a minor fraction of the microbiome (i.e. less than 16% of the microbial species) was specialised in polysaccharides’ degradation, while all the remaining planktonic species were involved in different steps of the anaerobic digestion food chain. All the firmly attached to grass PGs belong to *Firmicutes* and *Bacteroidetes*/*Chlorobi* phyla (Fig. 3) confirming previous findings based on 16S rRNA clone library analysis (17). The identification of a large number of *Firmicutes* was expected, since they are one of the dominant phyla in proficient plant biomass-degrading ecosystems like the rumen (22). At a lower taxonomic level, these PGs were assigned to the orders *Clostridiales* and *Thermoanaerobacterales*. On the contrary, the identification of species belonging to the phylum *Bacteroidetes* in the firmly attached to grass sample is intriguing because uncultured species assigned to this taxonomic group have been recently identified in the rumen and also in other cellulose degrading microbial communities (19, 21). Additionally, the numerical predominance of uncultured *Bacteroidetes* species in lignocellulosic-degrading environments suggests a relevant contribution of these microbes to polysaccharide degradation (22, 29, 30). The genomic properties of the extracted PGs were investigated to determine the rationale beyond their spatial localisation, emphasising on the microbes having a putative role in lignocellulose decomposition. The result from COG analysis showed that the “firmly attached PGs” were more enriched in “T-Signal transduction mechanisms” and “N-Cell motility”. Despite the fact that COG functional categories are very general, this result indicates that once grass particles are introduced into the anaerobic system, specific microbial species are able to perceive this signal, move towards the plant material and ultimately attach firmly onto it so as to efficiently metabolize and internalize carbohydrates. Previous studies demonstrated that chemotaxis and motility are widely diffused properties in lignocellulose degrading systems; different polysaccharides, such as cellobiose, cellotriose, D-glucose, xylobiose and D-xylose, can serve as chemoattractants for cellulolytic bacteria that migrate toward plant materials and decompose the organic substances (31, 32). Moreover, the outcomes from the dbCAN database demonstrated that the PGs that were firmly attached to the grass particles had more genes encoding cellulases, while the planktonic PGs had more debranching enzymes. This finding suggests a co-operative function between the two microbial clusters. The exoglucanases of PGs present in the liquid fraction or in *Eubacteriaceae* species (which were similarly enriched in liquid and grass fractions), such as *Eubacteriaceae* sp. DTU376, could prevent a potential accumulation of tetrasaccharides and cellobiose. In other case, the accumulation of these compounds can in turn inhibit the cellulase activity of the firmly attached microbial species. In a previously studied thermophilic cellulose methanation consortium, a similar co-operative mechanism was found to be performed by *Spirochaetales* or *Thermotogales* with *Clostridiales* (19). An interesting finding was related with the mechanisms that are followed by the different microbial species to bind and degrade lignocellulose. As reported in the results section, numerous cohesin-dockerin pairs were found in *Clostridiaceae* sp. DTU079 and *Clostridia sp.* DTU196. Cohesin modules function as the main building blocks of cellulosomal scaffoldin, while dockerin modules anchor the catalytic enzymes to the scaffoldin or can be present in the C-terminus of the same proteins (29, 33). The high efficiency of cellulosome in binding the bacterial cell to the plant polysaccharides explains why the PGs posing this feature are highly enriched in the fraction that was firmly attached to the plant biomass.

On the contrary, the mechanism of polysaccharide utilization in the *Bacteroidetes* sp. DTU139 is completely different from that present in *Clostridiales*. An initial distinct change is related with the absence of cohesin and dockerin domains in *Bacteroidetes*. The enzymes catalyzing the conversion of cellulose in uncultured *Bacteroidetes* are not arranged in a cellulosome (21), despite a previously reported cellulosomal scaffoldin gene in *Bacteroides cellulosolvens* (34). Considering the model of starch utilization system (Sus) of *Bacteroides thetaiotamicron* (35) we can depict a polysaccharide utilization locus, where the cellulose is degraded to cellobiose by the “glycosyl hydrolase family 9-containing” enzymes, it is transported to the periplasmic space by the disaccharide transporter, cleaved to glucose by the cellobiose phosphorylase and finally transported to the cytoplasm (Fig. 5b). It can be reasonably assumed that the outer membrane receptor protein and the outer membrane sensor are involved in the regulation of the other genes present in this locus.

This model extends the role of *Bacteroidetes* in polysaccharides degradation beyond the “grass-degrading” environments, such as rumen, suggesting a more general involvement in anaerobic digestion systems. Our findings further show that important multifunctional enzymes, such as the cellulosome, can be remarkably different in species both due to the number and due to the roles of the subunits involved. In the present study, the metagenomic binning integrated with bioinformatics strategies for gene annotation represents an important tool to identify and investigate microbial species participating in polysaccharides degradation. This approach proves to be particularly useful to explore the unculturable species present in the anaerobic digestion microbial community, which are impossible to investigate by classical genomic approaches. An accurate genome annotation elucidated the diverse metabolic strategies, which are followed by specific microbial species to bind and degrade the plant material, providing to other community members the sub-products for methane production. Moreover, the genome sequence and the functional annotation of 25 PGs representing the microbial fraction firmly attached to the plant material, is a fundamental element towards the comprehension of the mechanism for polysaccharide utilization in biogas reactors. In particular, the three draft genomes that were adherent to the grass are perfect candidates for further biotechnological applications in cellulose degradation, such as production of platform chemicals (e.g. 3-Hydroxypropionic acid, xylitol etc.) or sustainable alternative biofuels (e.g. biobutanol, aviation fuels etc.).

## Materials and methods

### Biogas reactors’ configuration, DNA extraction and metagenome sequencing

Samples were collected during the steady state periods (i.e. stable biogas production with a daily variation of lower than 10% for at least 10 days) of two biogas reactors as previously described (36). The first reactor was co-digesting pig manure together with ensiled meadow grass, while the second reactor was treating only pig manure (mono-digestion process). Three replicate samples were collected from each reactor; genomic DNA extraction and sequencing were performed independently for each replicate to allow statistical evaluation of the population genome abundance in the different fractions. For the samples collected from the co-digesting reactor, approximately 3 g of grass were separated from the liquid fraction. Plant fibers present in the liquid fraction (i.e. sample denoted as “LG”) were removed by driving the sample through a 40-μm nylon cell strainer filter and by centrifugation at 500 g for 15 min. After this step, microbial cells were recovered with a centrifugation at 10000 g for 20 min at 4°C. The grass obtained after the separation of the liquid material was processed in order to recover the firmly attached microbes (i.e. sample denoted as “FG”) as previously described (37). The grass was washed with Basal Anaerobic (BA) medium (38) to remove loosely attached microbial cells. After this step, the firmly attached microbes were stripped off by incubating the grass at 4°C in anaerobic conditions with BA medium containing 0.15% Tween-80 and, subsequently, homogenizing the grass with a VWR VDI-12 homogenizer, 2 x 2 min bursts (VWR, Radnor, PA, USA). The fragmented plant material remaining after homogenization was removed with a mild centrifugation step (500 g for 15 min) and adherent cells remaining in the supernatant were recovered by centrifugation at 10000 g for 20 min, at 4°C. Samples collected from the reactor performing the mono-digestion (i.e. sample denoted as “PM”) were processed similarly to the liquid fraction collected from the co-digestion process.

Genomic DNA from each sample was extracted using PowerSoil^^®^^ DNA Isolation Kit protocol (MO BIO laboratories, Carlsbad, CA, USA). The quality and concentration of extracted DNA was determined using NanoDrop (ThermoFisher Scientific, Waltham, MA, USA) and Qubit fluorometer (Life Technologies, Carlsbad, CA, USA), respectively. Integrity of the genomic DNA was evaluated with agarose gel electrophoresis. The sequencing was performed using the Nextera DNA Library Preparation Kit (Illumina, San Diego, CA, USA). Libraries were paired-end sequenced (2 x 150 bp) by NextSeq 500 sequencer (Illumina, San Diego, CA, USA) using the mid output kit. Sequencing data can be found at Sequence Read Archive with identifier SRP074882, BioProject PRJNA319008.

### De novo metagenome assembly and binning process

*De-novo* assembly was performed with CLC Genomics workbench software v 5.1 (CLC Bio, Aarhus, DK) using CLC’s *de novo* assembly algorithm (kmer 63, bubble size 60 and minimum scaffold length 500 bp). The binning process used to extract the population genomes from the assembly was previously reported by Campanaro et al., (26). Specific modifications and detailed parameters followed in the current study are provided in the Supplemental Information S1. It was previously demonstrated (39) that in the binning methods based on co-abundance clustering, the specificity of the genome extraction process suddenly reaches the maximum level when more than 15 different samples are used. For this reason, additional samples from previous metagenomic studies were included in the analysis (27, 26, 12). The additional samples were only used to assist the alignment of the scaffolds and to increase the total number of coverage conditions examined. The completeness and the genome contamination of each PG were determined by considering the total number of essential genes and the number of duplicated genes in all the examined trials.

### Taxonomic assignment of the population genomes

Taxonomic assignment of the PGs was performed with PhyloPhlAn software using 400 broadly conserved proteins used to extract phylogenetic signal (40). For 17 PGs, whose assignment was at low or incomplete confidence with PhyloPhlAn, the result was verified and/or corrected using Phylopythia classifier (41).

Identification of the common PGs between different metagenomic assemblies was performed by comparing the genomic sequences of the recovered PGs of the present study with those identified in previous metagenomic binning analyses related to biogas reactors and with the 60,585 genomes deposited in the NCBI “Microbial Genomes Resources” database (March 2016). The comparison was based on the calculation of the Average Nucleotide Identity (ANI) value (42) on the protein-encoding nucleotide sequences as previously described (27). Initially, a database with the nucleotide sequences of the PGs belonging to previous binning processes was obtained (27, 26). Similarity search was performed using BLASTn with all the nucleotide sequences of the genes identified for each PG, by using 1e^−5^ as minimum threshold. An in-house developed script, called ‘‘ANI_calculator_CS.pl” (https://biogasmicrobiome.com/), determined in the BLASTN output the number of sequences having a match in the database and the ANI value for each PG. As previously defined, two genomes were considered as belonging to the same species if at least 50% of the genes found a match and if the ANI was higher than 95% (42). Scaffolds encoding the 16S rRNA genes were identified using sequence similarity search on the Greengenes database, as previously proposed (43). Taxonomical assignment of the identified 16S rRNA genes was determined using RDP classifier (44) with confidence threshold set at 0.8.

### Quantitative PCR

The RT – PCRs were performed with the CFX96 Real-Time System (Bio-Rad, Hercules, CA). The PCR was performed using the iTaq SYBR Green Supermix (Bio-Rad, Hercules, CA) using the following conditions: pre-heating, 5 min at 95°C; cycling, 40 cycles of 15 s at 95°C, 15 s at 50°C and 60 s at 72°C. Results were expressed in terms of fold difference between the two fractions examined (firmly attached to grass and liquid) as determined by the difference in the Ct. Universal primers used for RT-PCR were 27F, 1492R for Bacteria and 109F, 1492R for Archaea.

### Functional and statistical analysis

Gene finding was performed in the assembled scaffolds using Prodigal software set in metagenomic mode (45). All the proteins predicted from the assembly were annotated using reverse position-specific BLAST algorithm (RPSBLAST of NCBI BLAST +) using different databases depending on annotation; for Clusters of Orthologous Groups of proteins (COG) the “COG only” RPSBLAST database was used (46), while for conserved protein domains prediction (Pfam) the Pfam RPSBLAST database was employed (47). Blast results were filtered with e-value lower than 1e^−5^ and, additionally only the best match was considered for COG. KEGG annotation was performed with the software GhostKOALA (48). Annotation of carbohydrate-active enzymes was performed using the software dbCAN (49). Functional annotation was considered only for the PGs having an estimated completeness higher than 50% as high number of missing genes in incomplete PGs would affect the evaluation of the results. For KEGG, COG and dbCAN annotation, the results obtained for all the predicted proteins were separated and assigned to the single PGs using in-house developed perl scripts. After the assignment process, the number of proteins belonging to each COG category (t-test, two tails, p value < 0.05), KEGG pathway and carbohydrate-active module were calculated for each PG. The classification of the glycosyl hydrolases in different functional categories “cellulases”, “endohemicellulases”, “debranching enzymes”, “oligosaccharide-degrading enzymes” and “other glycosyl hydrolases” was obtained from Hollister and coll. (50). The scaffolds of each PG were uploaded to the RAST server and automatically re-annotated using SEED and the RAST gene caller (51). Annotation results were downloaded and the number of genes present in each first and second level SEED category was determined using in-house developed perl scripts. A random sampling process was repeated 1000 times with a Perl script implementing the Perl “rand()” function (52) to determine if the PGs are enriched in specific functional categories. The same number of proteins encoded in each PG were randomly collected from the entire assembly and assigned to each functional category. Finally, the fraction of random samples in which the number of genes assigned to a specific functional category was equal or higher than N (where N is the number of genes assigned to the same functional category in the PG) was determined. The enrichment in a specific functional category was considered as significant when the fraction was lower than the significance level (α = 0.05).

### Spatial assignment of the population genomes

The classification of the PGs as firmly attached to grass, planktonic and that with similar enrichment in both fractions (i.e. grass and liquid) was obtained by comparing the coverage of the PGs in the replicate samples “FG” and “LG”. Those enriched more than 2 fold in “FG” were considered as “firmly attached”, those enriched more than 2 fold in the liquid fraction were considered and “planktonic” and the remaining PGs were assigned to the group with intermediate behaviour. Consistency of the results among the three replicates was determined using two tails t-test and selecting PGs with p-value lower than 0.05.

## Acknowledgement

We acknowledge financial support from Energistyrelsen under the programme EUDP, project 64013-0159 ‘‘New technology for an efficient utilization of meadow grass in biogas reactor’’. Illumina sequencing was performed at the Ramaciotti Centre for Genomics (Sydney, Australia).

